# Salivary microbiome diversity is associated with oral health and disease

**DOI:** 10.64898/2026.01.27.702135

**Authors:** David L. Lin, Madisyn D. Augustine, David M. Ojcius

## Abstract

The oral microbiome is a complex community of bacteria, fungi, and viruses that inhabit the oral cavity. Microbes of the oral microbiome are implicated in health and disease. We collected 220 unstimulated saliva samples from patients with periodontal disease and varying degrees of dental caries, as well as from subjects with no signs of oral disease. Metagenomic analysis of saliva revealed significantly higher abundance of periodontal pathogens in people with gum disease, and significantly higher abundance of cariogenic species in people with dental caries. We also found that salivary microbiome diversity was significantly higher in people with periodontal disease, but not in those with only caries. Furthermore, oral microbiome diversity is affected by oral hygiene habits such as flossing frequency, but not brushing frequency. Clustering and differential analysis allowed us to identify specific commensal species, such as *Prevotella pallens and Veillonella atypica*, which are significantly higher in patients without oral disease. Clustering further suggested that oral microbiome diversity may contribute to disease risk. These results suggest that oral hygiene behaviors influence the oral microbiome, and modulation of the oral microbiome could prevent or reduce the incidence and severity of oral disease.

## Introduction

Oral diseases such as periodontal disease and dental caries are two of the most prevalent oral conditions in the world, and periodontal disease is one of the most common inflammatory diseases in humans. Over 50% of adults in the US have some form of gum disease, and 25% of adults have untreated tooth decay ^1^. Decades of research have shown causal relationships between microbes living in the oral cavity and both oral and non-oral systemic disease ^2^. Despite these associations, diseases are still commonly diagnosed only after the progression of visible symptoms, and oral disease is still treated using traditional techniques that involve the physical removal of oral biofilm.

The oral microbiome is a community of bacteria, viruses, and fungi that play a critical role in maintaining oral health and homeostasis. Over 700 different species of bacteria have been identified in the oral microbiome, making it the second most diverse microbiome in the human body after the gut microbiome. Similar to the gut, dysbiosis in the oral microbiome has been linked to many oral and systemic diseases, including Alzheimer’s disease, colorectal cancer, and diabetes^3^. However, the mechanisms linking oral microbes to systemic disease remain unclear. By contrast, the oral microbiome is a critical component in the development and incidence of oral disease, with known mechanisms driving health and disease.

Key species in the oral microbiome have been shown to play causal roles in oral disease. For example, *Streptococcus mutans* and *Streptococcus sobrinus* are well-known cariogenic species capable of driving tooth decay ^4^. Similarly, specific anaerobic periodontal pathogens, such as *Porphyromonas gingivalis* and *Tannerella forsythia*, cause chronic gum inflammation, which can eventually lead to bone loss and periodontal disease ^5^. Despite decades of research on periodontal disease, the mechanisms that drive disease are still unclear ^6^. However, community-level interactions between oral microbes have been shown to play important roles in oral microbiome homeostasis and dysbiosis.

Interactions between oral microbes are important in maintaining health. For example, *Streptococcus sanguinis* antagonizes the growth of pathogenic *S. mutans* and increases the local pH in biofilms ^7^. Numerous species have been associated with health, including Streptococci, Neisseria, and Haemophilus species ^8^. Some in vitro studies have shown that these commensals can directly inhibit the growth of periodontal pathogens, although the mechanisms of inhibition are not always known ^9^.

In this study, we used shotgun metagenomic sequencing to profile the salivary microbiome in disease and health of 220 people. We found that oral disease was associated with distinct differences in the salivary microbiome, and oral hygiene habits were also associated with apparent shifts in the community. We demonstrate that oral microbiome diversity is correlated with periodontal disease but not dental caries, suggesting diversity may be a predictor of gum health. We were able to identify a number of species associated with health through hierarchical clustering. An improved understanding of the oral microbiome and its role in health and disease could lay a foundation for the development of novel therapeutics that modulate the oral microbiome to prevent and treat oral disease.

## Methods

### Study population and sample collection

Participants were recruited from patients at the dental clinic of the University of the Pacific Arthur A. Dugoni School of Dentistry. A total of 220 participants were enrolled between 2021 and 2022. The age of participants ranged from 17-60 years old. After signing informed consent, each participant completed a demographics questionnaire related to oral health, oral hygiene, overall health, and demographics. Unstimulated saliva samples were collected before any dental treatment was performed. Sample collection was approved by the Institutional Review Board of the University of the Pacific.

### Clinical assessment

153 of the 220 participants underwent clinical assessment to diagnose gum disease or dental caries. Periodontitis was defined as interdental clinical attachment loss of > 1mm and classified according to the American Academy of Periodontology (AAP) guidelines ^10^. Caries were visually detected and classified according to the International Caries Detection and Assessment System (ICDAS) ^11^. Those lesions with ICDAS codes of “5” or “6” were considered to be cavities in this study.

### Library Preparation and Sequencing

Zymo Research DNA/RNA Shield SafeCollect Saliva Collection Kits were used to collect and store saliva samples according to the manufacturer’s instructions. Sample lysis and purification were performed according to the Zymobiomics DNA/RNA lysis and purification protocol with some modifications: 1 ml of saliva was lysed via bead beating with a tabletop vortex per the manufacturer’s instructions. Insoluble fractions were removed by centrifugation, and the soluble fraction was subjected to proteinase K digestion prior to nucleic acid purification. DNA was purified using Zymobiomics Magbead purification per manufacturer’s instructions. Sequencing library preparation was performed according to Illumina DNA Prep by tagmentation per manufacturer instructions with reduced PCR amplification to 11 cycles. DNA quality and quantity were measured using Qubit fluorometric quantification and a bioanalyzer. Libraries were sequenced (2×150bp) on a NovaSeq to an average of 13 million reads per FASTQ.

### Bioinformatics

Reads were pre-processed by removing adaptors with low-quality bases. Reads were quality trimmed using cutadapt. Fastqs were then downsampled to 5M paired-end reads to prevent potential downstream bias. Human reads were removed by aligning reads to the human reference genome GRCh38 using Bowtie2. To characterize the microbial composition of the remaining non-human reads, paired-end reads were aligned using MIDAS to bacterial marker genes ^12^. Taxonomic count tables and functional counts were analyzed using custom R scripts for clustering and statistical analysis. Maaslin2 was used to discover associations between phenotypes and microbial abundance.

## Results

### Clinical summary

Out of 153 participants who underwent clinical assessments, 50 were clinically healthy, free of tooth decay or gum disease, while 34 were diagnosed with periodontal disease alone, 31 with dental caries alone, and 38 with both. The remaining 67 samples in this study did not have accompanying clinical assessments, although some participants submitted survey responses about hygiene behavior and demographics.

### Bioinformatic summary

We sequenced 220 saliva samples with an average depth of 13 million reads. After downsampling, host-depletion, and microbial mapping, we detected a median of 117.5 bacterial species per sample and a total of 527 bacterial strains representing 328 total species observed. Strains were collapsed into species by summing the relative abundance of multiple strains that contribute to a single species. A total of 171 species were present in more than 20% of all samples.

### Salivary microbiome archetypes reveal associations with disease

To better understand community composition, we used hierarchical clustering to group salivary microbiomes, which we termed oral microbiome archetypes. We found that archetypes were grouped loosely by the most abundant species of the community, but were also associated with disease status (Figure 1). We grouped archetypes into five distinct clusters, each characterized by common core microbial species in that group. These clusters also revealed that some community types were more associated with disease (Table 1). The overall prevalence of disease was 47% and 45% for periodontitis and dental caries, respectively. Notably, Cluster 5 had the overall highest rates of disease, with 93% diagnosed with periodontitis and 53.3% with caries. Cluster 3 had the lowest rates of disease, with only 34.6% with periodontitis, and 38.4% with caries.

**Table 1.** Clusters by disease prevalence. The prevalence of disease in each cluster, frequency of each cluster, and total number of samples associated with each are shown.

|  | Periodontal disease prevalence (%) | Caries prevalence (%) | Overall cluster frequency (%) | Samples in cluster |
| --- | --- | --- | --- | --- |
| Cluster 1 | 46.70 | 46.70 | 10.45 | 23 |
| Cluster 2 | 44.00 | 28.00 | 33.64 | 74 |
| Cluster 3 | 34.60 | 38.50 | 19.09 | 42 |
| Cluster 4 | 46.50 | 62.80 | 28.18 | 62 |
| Cluster 5 | 93.30 | 53.30 | 8.64 | 19 |
| Overall | 47.06 | 45.10 | 100.00 | 220 |

**Figure 1.**
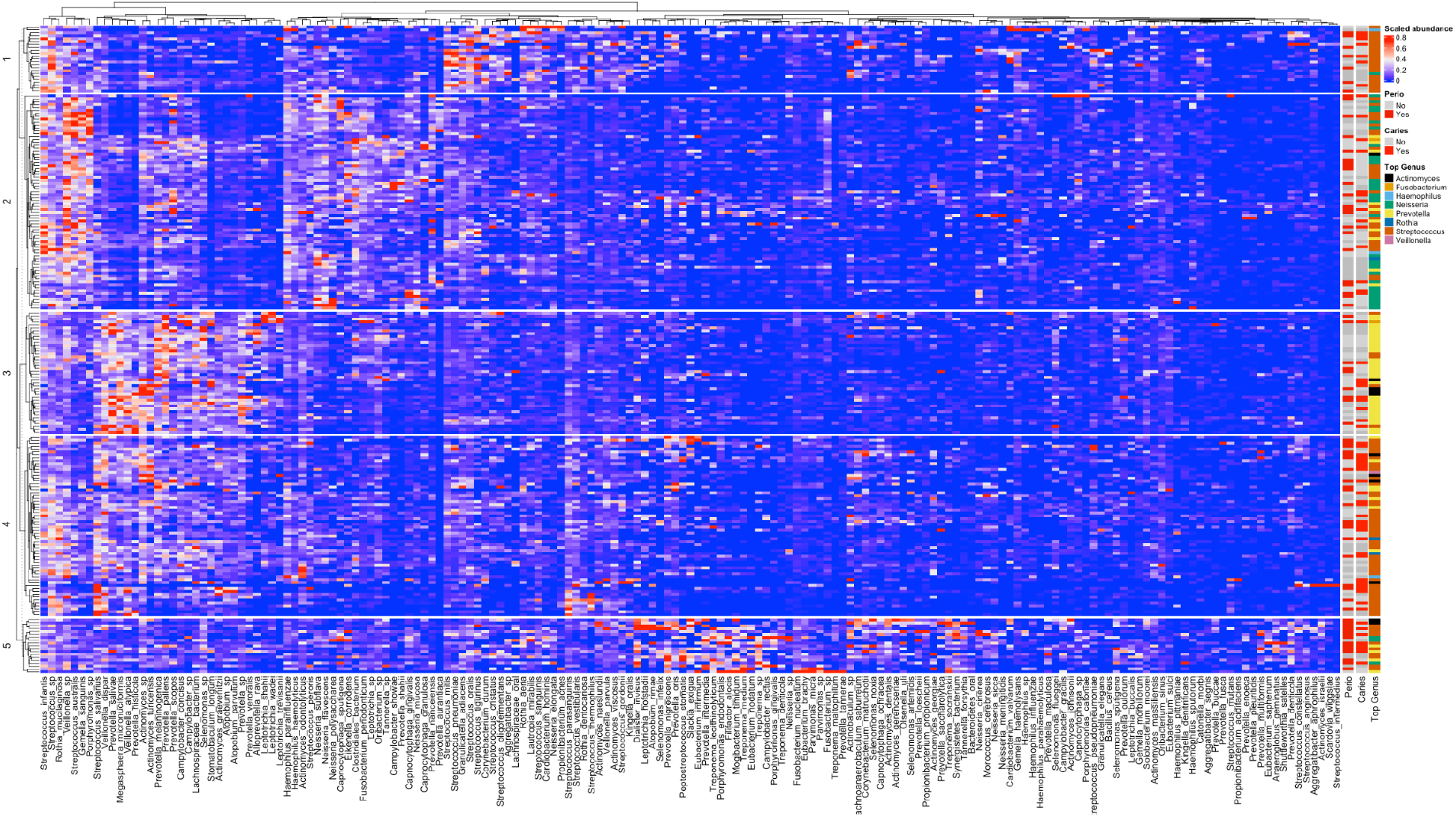
Heatmap of all salivary microbiomes for the species that were detected in at least 10% of all samples. The y-axis represents individual samples and the x-axis represents different species. Relative abundance of each species is scaled from 0 to 1. Hierarchical clustering differentiates between groups of samples that are most similar to each other. The genus of the most abundant species in a sample is indicated on the right annotation, along with the disease status of the participant from where the sample originated. Samples without clinical metadata are indicated in darker grey. Species are organized across the x-axis by hierarchical clustering.

### Community dynamics in health and disease

To better understand how Cluster 3 may be associated with better oral health, we used multivariable association analysis to identify community members that are most prevalent and abundant in the cluster. Unsurprisingly, we found that canonical periodontal pathogens, such as *Eubacterium brachy, Tannerella forsythia, Treponema* species, *Filifactor alocis, Fusobacterium nucleatum, Parvimonas micra*, and *Prevotella intermedia* were significantly higher in Cluster 5 compared to other clusters (Figure 2A-C). Additionally, *Mogibacterium timidum* and *Anaeroglobus geminatus* were enriched in Cluster 5, which have recently been identified as periodontal pathogens (Figure 2A-C) ^13^. We also found a higher prevalence of caries-associated species *S. mutans* and *Propionibacterium acidifaciens* in Cluster 5, further corroborating the high rates of disease in this archetype (Figure 2A).

**Figure 2.**
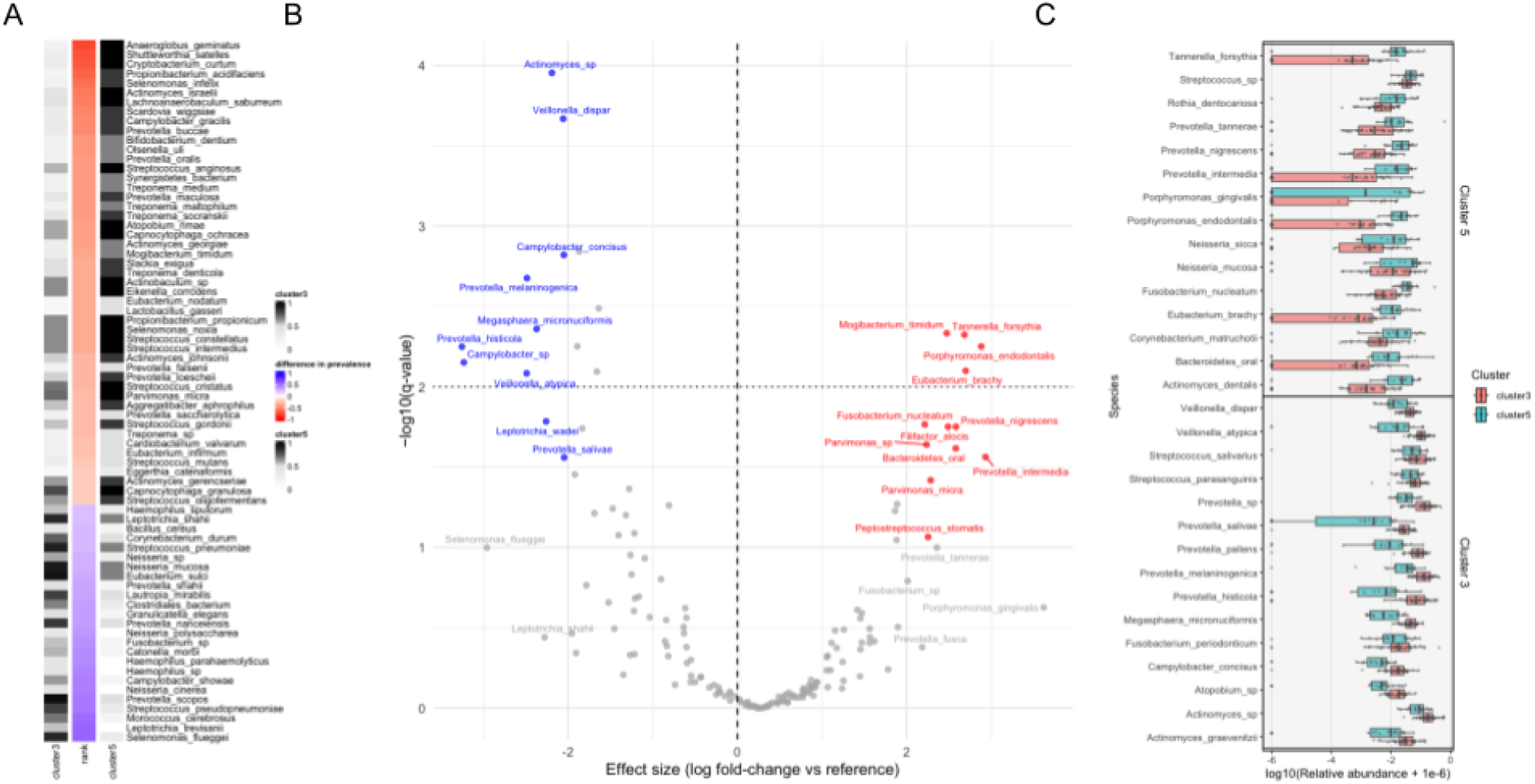
Differentially abundant species in cluster 3. In A, species most differentially prevalent between clusters 3 and 5. The total prevalence for each species is in grayscale, with darker black being higher prevalence. The difference in prevalence between clusters is color scaled from blue to red. In B, a volcano plot of differential species between clusters 3 and 5. Species significantly associated with cluster 5 are in red, and cluster 3 in blue. Species in gray are associated with either cluster, but with q values less than 0.1. In C, the relative abundance of the top 15 most differentially abundant species between the two clusters are plotted. Each point represents an individual patient, and the x-axis is log10 of the relative abundance + 1 x 10^6^.

Alternatively, we found that *Actinomyces, Atopobium, Leptotrichia, Prevotella*, and *Veillonella* species were enriched in Cluster 3 (Figure 2A). Surprisingly, we also found that Cluster 3 had the lowest abundance of *Streptococcus* species, such as *Streptococcus mitis, Streptococcus gordonii*, and *Streptococcus sanguinis*, which are species often associated with oral symbiosis^14^. We found that Cluster 1 had the highest abundance of Streptococci, with rates of disease equivalent to the overall prevalence of disease. These data suggest that abundance of commensal Streptococci is a poor predictor of disease, whereas the abundance of *Veillonella* and *Prevotella* species may be indicators of symbiosis and oral health. However, some *Streptococcus* species were enriched in Cluster 3, such as *S. salivarius* and *S. parasanguinis*. Cluster 3 had the highest abundances of *Veillonella atypica* and *V. dispar*, two species often associated with oral health.

We found that *Prevotella* species were particularly disparate between clusters. *P. tannerae, nigrescens*, and *P. intermedia* were associated with Cluster 5, while *P. pallens, P. melaninogenica*, and *P. histicola* were associated with Cluster 3. This data suggests that a shift in the abundance of *Prevotella* species may be a hallmark of dysbiosis.

### Oral microbiome clusters reveal associations between disease and diversity

Next, we measured diversity by Shannon index, and found a noteworthy difference, where Cluster 5 exhibited a significantly higher diversity compared to all other clusters (Figure 3A). We also found that diversity was significantly higher in people with periodontal disease compared to people without periodontal disease (Figure 3B). By contrast, we found that diversity was unchanged between people with dental caries compared to those without caries (Figure 3C). These data suggest that increased oral microbiome diversity may be a risk factor for periodontal disease, but not dental caries.

**Figure 3.**
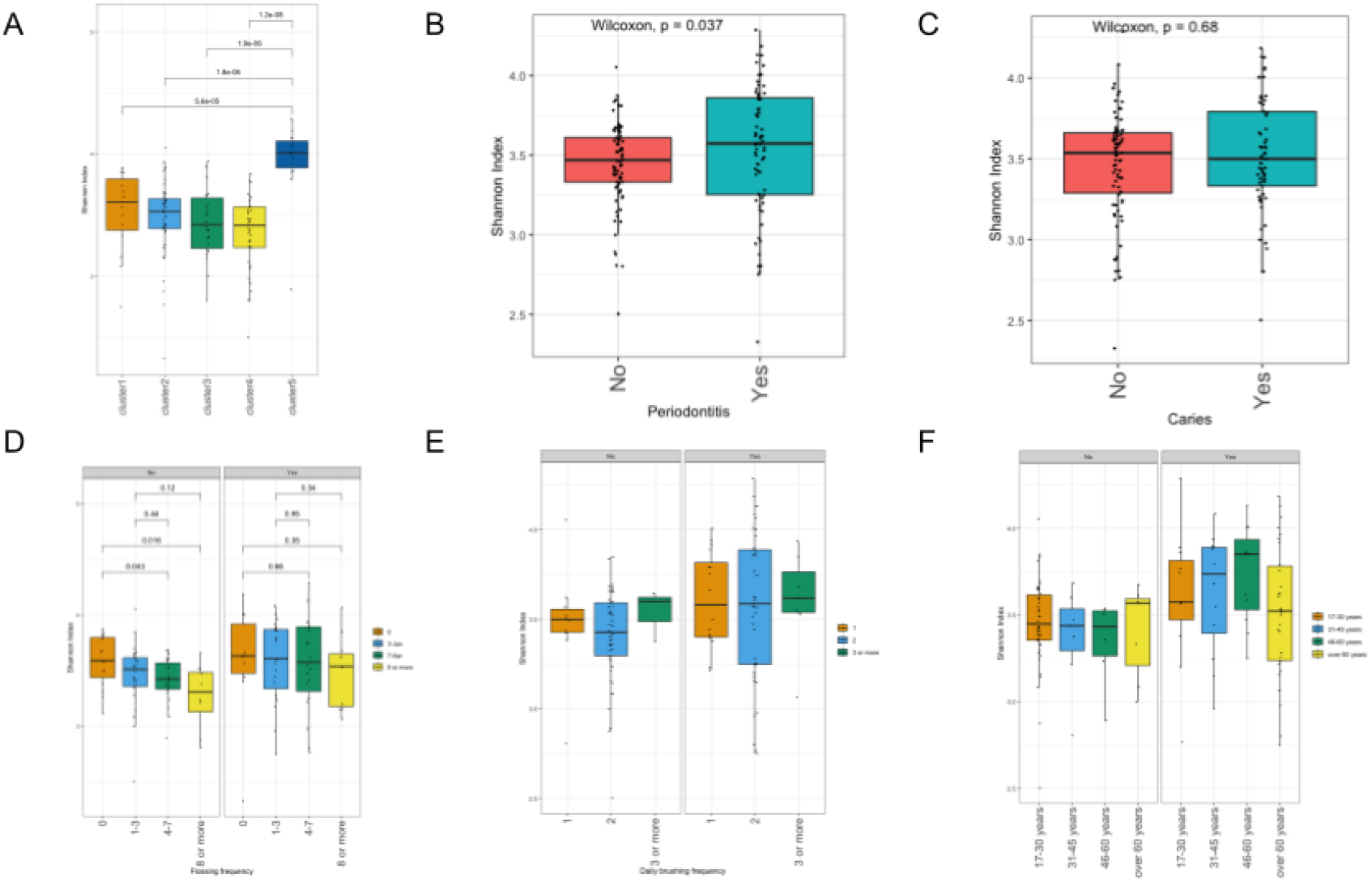
Oral disease and hygiene habits affect oral microbiome diversity. The y-axis represents diversity, which was measured by shannon index, and each point represents a different individual, with boxplots showing the interquartile range and median. Points are jittered for visualization. In A, diversity is grouped by cluster. In B and C, diversity is grouped by disease status as indicated on the x-axis. In D, E, and F, the groups “No” and “Yes” indicate either healthy gums or periodontal disease. The x-axis represents weekly flossing frequency, daily brushing frequency, and age respectively. Where indicated, statistics were calculated by mann-whitney u test between the indicated pairs.

### Oral microbiome diversity is affected by oral hygiene routines

We next assessed how oral hygiene routines affect the oral microbiome. Few studies have been done in vivo with respect to the effect of brushing and flossing on the oral microbiome. Using self-reported oral hygiene surveys, we found that flossing frequency was highly correlated with oral microbiome diversity, which decreased with increased flossing frequency (Figure 3C). By contrast, brushing frequency was not correlated with oral microbiome diversity regardless of disease status (Figure 3D). Additionally, we looked at whether age was a major factor in oral microbiome diversity, and found that for healthy people, age generally did not affect diversity (Figure 3E). Flossing frequency was not associated with any cluster (Figure S1). Together, these results and the increased diversity observed in periodontal disease suggest that for people without periodontal disease, flossing may reduce the risk of developing periodontal disease by reducing the oral microbiome diversity. However, flossing frequency does not reduce diversity for people who already have irreversible periodontal disease.

## Discussion

For decades, research has shown causal relationships between oral microbes and oral and systemic disease. Our results corroborate decades of research demonstrating an increased abundance of pathogenic species in the two most common oral diseases, gum disease and caries. For instance, differential analysis of shotgun metagenomic data identified red complex species *T. forsythia, P gingivalis, T. denticola* as increased in people diagnosed with periodontitis. Similarly, we found a significantly increased abundance of *S. mutans* and *P. denticola* in people with active caries ^15,16^. This data provides strong validation for shotgun metagenomics as a reliable approach to characterize the oral microbiome and oral disease.

One characteristic of microbiome dysbiosis is a general loss of commensal species ^17^. Hierarchical clustering of oral microbiomes revealed five distinct groups of microbiome types, grouped generally by the most abundant species in their community. These clusters revealed potentially protective species that are associated with health, such as *Prevotella pallens* and *P. histicola*, directly contrasting *Prevotella* species that may be involved in disease and dysbiosis such as *P. intermedia*. Through clustering, we could demonstrate that core microbiomes were significantly different between healthy people and those with periodontal disease and caries. While unsupervised clustering revealed distinct and robust clusters, these microbiome archetypes should be viewed as descriptive frameworks rather than definitive disease states.

These clusters also revealed that grouping was highly associated with periodontal disease and oral microbiome diversity. Typically, ecological diversity is a sign of the health of a community. In the gut, increased diversity is associated with ecological stability and health. However, our data suggest that, unexpectedly, increased diversity is associated with gum disease, similar to other recent studies^18,19^. We hypothesize that niches in the oral cavity are created through inflammatory-mediated tissue damage beneath the gumline, which fosters an environment that allows pathobionts to propagate unimpeded by normal commensal microbes. The expansion of these niches could lead to higher diversity and progression of irreversible periodontal disease. While this same principle could be applied to caries, we didn’t find any difference in diversity between people with active carious lesions and those without. It is important to note that salivary diversity is not equivalent to subgingival diversity, but shotgun metagenomic sequencing of saliva may be a useful tool to measure overall oral microbiome diversity.

We found a clear association between diversity, disease, and clusters. We were able to identify species associated with dysbiosis; however, their role in disease is still unclear. For example, we hypothesize that some *Prevotella* species may play a protective role, precluding colonization by pathogenic *Prevotella* species. Similarly, *Veillonella dispar* and *V. atypica* were strongly associated with oral health. This data may suggest that oral probiotics containing these protective species could improve outcomes for people with periodontal disease or prevent disease altogether.

Clinical studies have demonstrated the importance of flossing in maintaining oral health. This is the first study to associate the benefit of flossing with changes in the oral microbiome and oral health. We demonstrate that diversity decreases with increased flossing frequency, potentially revealing a mechanism by which flossing can reduce the risk of periodontal disease through modulating the oral microbiome. This conclusion is strengthened by the dose-dependency of flossing frequency and its impact on oral microbiome diversity. Additionally, scaling and root planing and antibiotic treatment as periodontal therapy reduces oral microbiome diversity, potentially describing one mechanism by which periodontal therapy improves prognosis^20^.

## Acknowledgements

We would like to thank the students at the University of the Pacific for assisting in patient recruitment and saliva collection. We are grateful for the support of staff at the University of the Pacific, including Dr. Des Gallagher and Harmony Matshik Dakafay. We would also like to thank Dr. Nader Nadershahi for encouraging faculty and students at Dugoni School of Dentistry to participate in this research.

## Ethics Statement

This study was approved by the Institutional Review Board at the University of the Pacific (IRB protocol #2020-87). Written informed consent was obtained from all participants.

## Author Contributions

David L. Lin (DLL) and David M. Ojcius (DMO) designed the study. DMO recruited students, managed sample collection and storage. Madisyn D. Augustine (MDA) and DLL performed wet lab experiments and bioinformatic analyses. DLL wrote the manuscript, which DMO and MDA critically revised. All authors signed off and approved the manuscript.

---

**S1.**
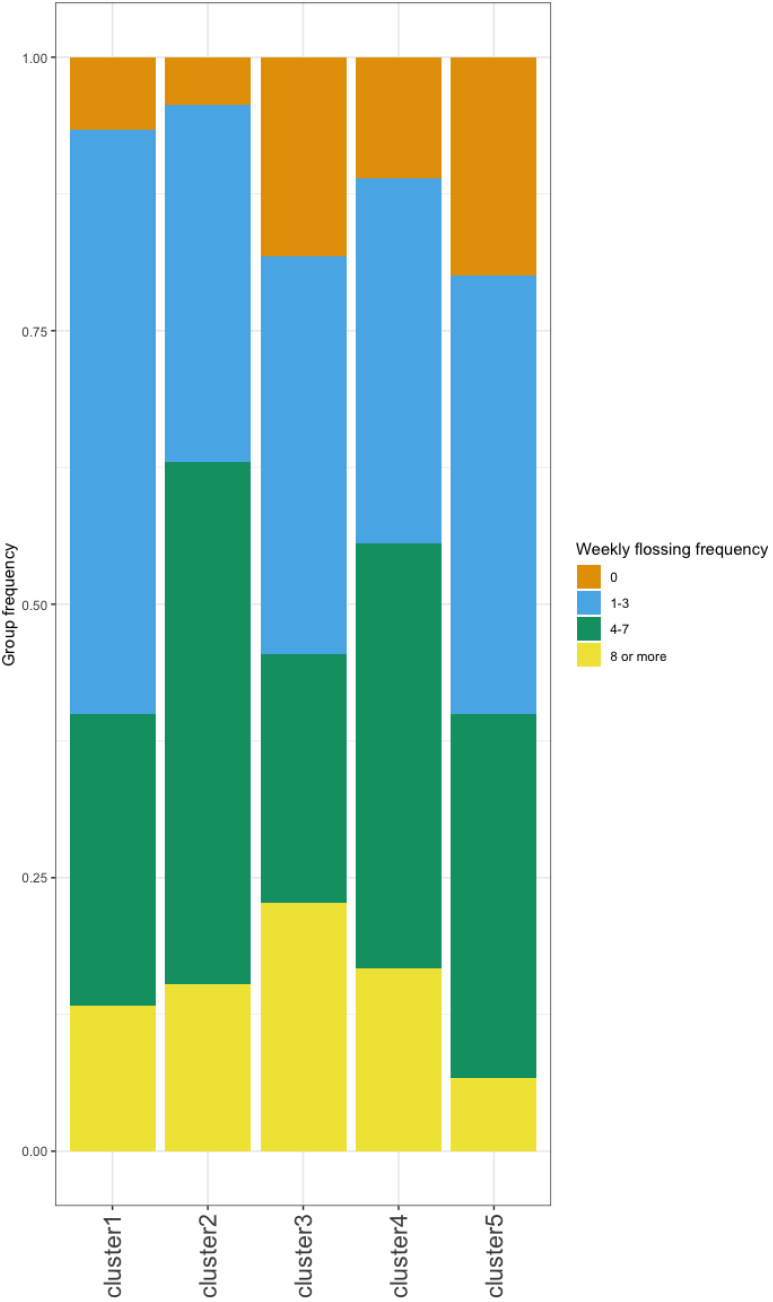
Distribution of flossing frequency by cluster. Each color is the fraction of people that floss at the indicated frequency. Orange is 0 times per week, blue is 1-3 times per week, green is 4-7 times per week, and yellow is 8 or more times per week.

